# Multivariate pattern analysis of MEG and EEG: a comparison of representational structure in time and space

**DOI:** 10.1101/095620

**Authors:** Radoslaw Martin Cichy, Dimitrios Pantazis

## Abstract

Multivariate pattern analysis of magnetoencephalography (MEG) and electroencephalography (EEG) data can reveal the rapid neural dynamics underlying cognition. However, MEG and EEG have systematic differences in sampling neural activity. This poses the question to which degree such measurement differences consistently bias the results of multivariate analysis applied to MEG and EEG activation patterns. To investigate, we conducted a concurrent MEG/EEG study while participants viewed images of everyday objects. We applied multivariate classification analyses to MEG and EEG data, and compared the resulting time courses to each other, and to fMRI data for an independent evaluation in space. We found that both MEG and EEG revealed the millisecond spatio-temporal dynamics of visual processing with largely equivalent results. Beyond yielding convergent results, we found that MEG and EEG also captured partly unique aspects of visual representations. Those unique components emerged earlier in time for MEG than for EEG. Identifying the sources of those unique components with fMRI, we found the locus for both MEG and EEG in high-level visual cortex, and in addition for MEG in early visual cortex. Together, our results show that multivariate analyses of MEG and EEG data offer a convergent and complimentary view on neural processing, and motivate the wider adoption of these methods in both MEG and EEG research.

## 2 Introduction

Multivariate pattern analysis of magnetoencephalography (MEG) and electroencephalography (EEG) data provide a fine-grained characterization of the temporal dynamics of neural activity. Recent research efforts have applied multivariate analyses, such as pattern classification and representational similarity analysis (RSA) (Kriegeskorte and Kievit, 2013a), in a rapidly expanding range of studies, demonstrating that MEG and EEG signals contain information about a diverse array of sensory and cognitive processes (e.g. see Groen et al., 2013; Cichy et al., 2014; Clarke et al., 2014; Isik et al., 2014; King and Dehaene, 2014; Kaneshiro et al., 2015; Kietzmann et al., 2016).

While in principle MEG and EEG signals arise from the same neuronal sources, typically postsynaptic currents from apical dendrites of pyramidal cells in cortex, there are consistent physical differences in the generated magnetic and electric fields (Cohen and Hosaka, 1976; Cohen and Cuffin, 1983; Hämäläinen et al., 1993) for several reasons. Radially-oriented sources are prominent in EEG but nearly silent in MEG, suggesting the existence of unique information coded in EEG signals. Further, the MEG and EEG spatial patterns of tangentially-oriented sources are 90^o^ relative to each other, leading to differential spatial sampling of neural activation. Also, EEG has higher sensitivity to deep sources than MEG. Unlike MEG, though, volume currents measured by EEG are deflected and smeared by the inhomogeneity of the tissues comprising the head.

These differences suggest that MEG and EEG are sensitive to partly common, and partly unique aspects of neural representations. This has been asserted by a large body of previous research encompassing theoretical argument as well as practical and experimental investigations mainly in the context of source localization and epilepsy research (e.g., Leahy et al., 1998; Henson et al., 2003; Sharon et al., 2007; Molins et al., 2008; Fokas, 2009). However, the relation of MEG and EEG in the context of multivariate analysis methods such as classification and RSA has not been investigated. Thus, two open questions remain: how comparable are the results of multivariate analyses when applied to either MEG or EEG, and to which extent do they resolve common or unique aspects of visual representations?

To address these open questions, we conducted an experiment with concurrent recording of MEG and EEG signals while participants viewed images of objects of different categories. We then applied equivalent multivariate pattern analyses to data from each modality and compared results in the time domain by 1) assessing the time courses with which objects and categories were discriminable by pattern classification, and 2) characterizing common vs. unique aspects of visual representations using representational similarity analysis. In space, we compared MEG and EEG by assessing the fusion of the temporally-informed MEG and EEG representations with spatially-informed fMRI representations using representational similarity analysis (Cichy et al., 2014, 2016c).

## 3 Methods

### 3.1 Participants

16 healthy human volunteers (7 female, age: mean ± s.d. = 24.1 ± 4.5 years, recruited from a participant pool at Massachusetts Institute of Technology) participated in the experiment. Written informed consent was obtained from all subjects. The study was approved by the local ethics committee (Institutional Review Board of the Massachusetts Institute of Technology) and conducted according to the principles of the Declaration of Helsinki.

### 3.2 Visual stimulus set and experimental design

The stimulus set consisted of 92 color photographs (Kiani et al., 2007; Kriegeskorte et al., 2008; Cichy et al., 2014, 2016b) of human and non-human faces and bodies, as well as natural and artificial objects isolated on a gray background (Fig. 1a). Participants viewed images presented at the center of the screen (4° visual angle) for 500ms and overlaid with a light gray fixation cross. Each participant completed 15 runs of 290 s duration each. Every image was presented twice in each run in random order, and the inter-trial interval (ITI) was set randomly to 1.0 or 1.1 s with equal probability. Participants were asked to maintain fixation and to press a button and blink their eyes in response to a paper clip image shown randomly every 3 to 5 trials (average 4). The paper clip image was not part of the 92 image set, and paper clip trials were excluded from further analysis.

**Fig 1.**
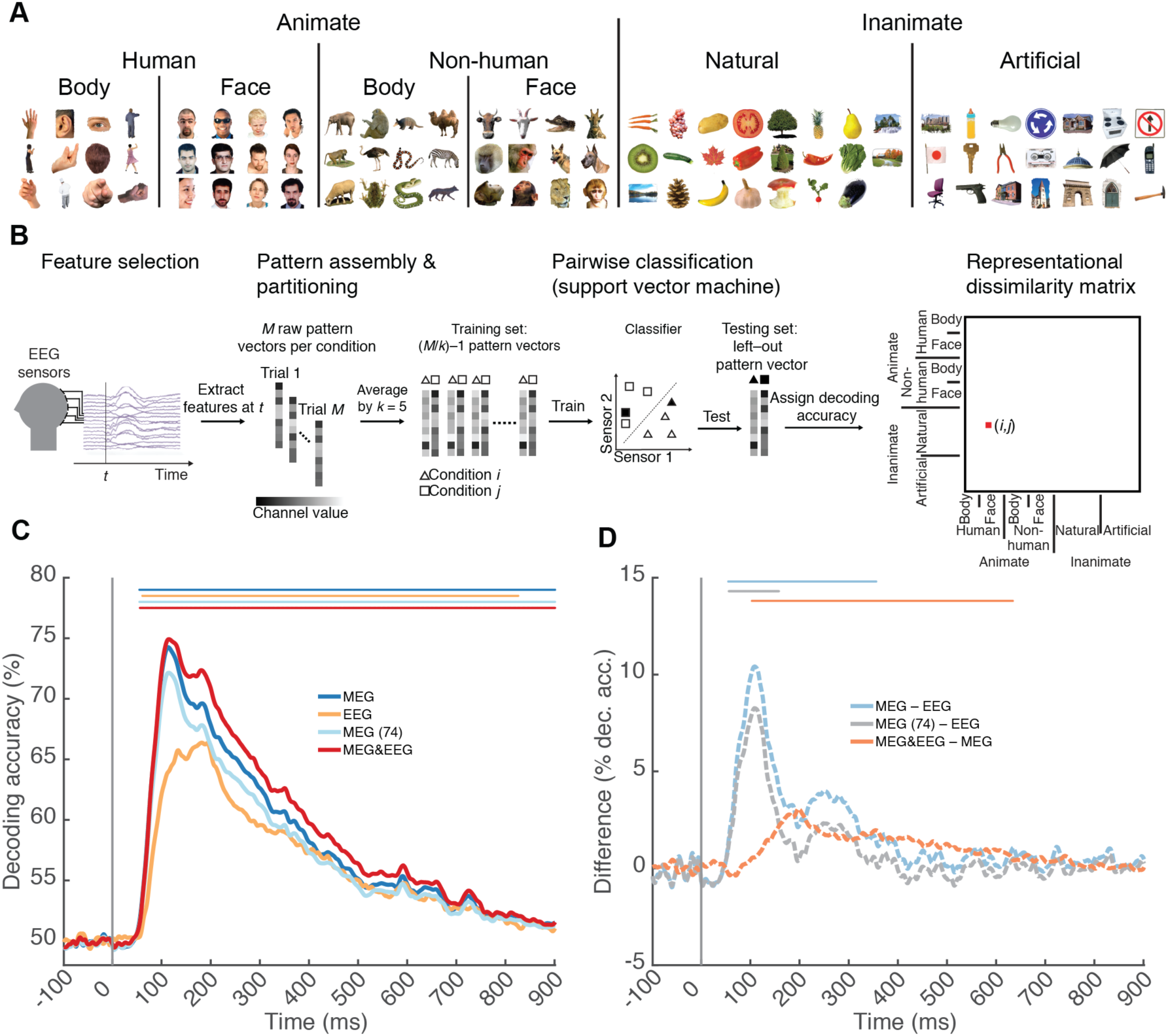
Classification of single images from EEG and MEG signals. **A**) The image set consisted of 92 silhouette images of everyday objects belonging to different categories. **B**) Multivariate pattern classification procedure, here shown for EEG data. **C**) Time courses of grand average decoding for different samplings of MEG and EEG sensors. **D**) Difference curves for the results shown in C. Lines above curves (same color-code) indicate significant time points (*n* = 15, cluster-defining threshold *P* < 0.05, corrected significance level *P* < 0.05 Bonferroni-corrected by number of plots for each subpanel, both two-sided). The gray vertical line indicates image onset. For equivalent results based on additional sensor samplings see Supplementary Fig. 2.

### 3.3 MEG and EEG acquisition and preprocessing

MEG and EEG signals were acquired simultaneously. We recorded MEG signals from 306 sensors (204 planar gradiometers, 102 magnetometers, Elekta Neuromag TRIUX, Elekta, Stockholm), and EEG signals from 74 sensors (custom-made cap with MEG compatible Ag/AgCl sensors; Easycap, Germany; sensor layout in Supplementary Fig. 1). Acquisition was continuous with a sampling rate of 1,000 Hz, and MEG/EEG data was filtered online between 0.03 and 330 Hz. Raw MEG data was preprocessed using Maxfilter software (Elekta, Stockholm) to perform noise reduction with spatiotemporal filters and head movement compensation (Taulu et al., 2004; Taulu and Simola, 2006). We applied default parameters (harmonic expansion origin in head frame = [0 0 40] mm; expansion limit for internal multipole base = 8; expansion limit for external multipole base = 3; bad sensors automatically excluded from harmonic expansions = 7 s.d. above average; temporal correlation limit = 0.98; buffer length = 10 s). Further preprocessing was carried out using Brainstorm (Tadel et al., 2011). In detail, we extracted the peri-stimulus MEG/EEG data of each trial from -100 to +900 ms with respect to stimulus onset, removed baseline mean, smoothed data with a 30Hz low-pass filter and divided the data of each sensor by the standard deviation of the pre-stimulus baseline signal of that sensor. This procedure yielded 30 preprocessed trials for each of the 92 images per participant.

### 3.4 Multivariate analysis

As the basis for multivariate pattern classification and subsequent comparison of MEG and EEG-based results in representational space, we sampled MEG and EEG data by sensors in four different ways: i) all 74 EEG sensors, ii) all 306 MEG sensors, iii) a random subset of 74 MEG sensors, thus equal to the number of EEG sensors, and iv) the combination of all 380 MEG and EEG sensors. In supplementary analyses we further report on other samplings of MEG and EEG data: i) 32 EEG sensors to determine whether basic EEG setups also enable multivariate analysis, ii) all magnetometers and iii) all gradiometers to investigate both types of MEG sensors separately, and iv) 74 magnetometers and v) 74 gradiometers to equate the number of sensors to EEG. We use the labels *MEG, EEG, MEG&EEG* for the full sensor arrays (306 MEG sensors, 74 EEG sensors, 380 M/EEG sensors), and provide the number of sensors in brackets only for reduced data sets, e.g. *MEG (74), EEG (32),* etc. All reduced data sets were constructed with equiprobable random samplings of the corresponding full sensor arrays (see next section).

Note that while we report results based on the four main sensor samplings in the main manuscript, for clarity we do not reference supplementary figures and tables with supplementary sensor samplings. These results are referenced in the main figure captions and tables, since they share the same formatting structure.

#### 3.4.1 Time-resolved single image classification

We first determined the time course with which single experimental conditions, i.e. images, are discriminated by MEG and EEG activation patterns (Fig. 1B).

Discrimination was assessed using linear support vector machine (SVM) classification (Müller et al., 2001), as implemented in the libsvm software (Chang and Lin, 2011) with a fixed regularization parameter C = 1. The classification approach was time-resolved, with pattern vectors created from MEG and EEG sensor measurements separately for every millisecond. In particular, for each time point *t* (from −100 to +900 ms in 1 ms steps), condition-specific sensor activation values for each trial (*M* = 30) were concatenated to pattern vectors, resulting in 30 raw pattern vectors. To reduce computational load and improve the signal-to-noise ratio, we sub-averaged the *M* vectors in groups of *k* = 5 with random assignment, obtaining *M/k* = 6 averaged pattern vectors. For all pair-wise combinations of conditions, we trained and tested the SVM classifier on the averaged pattern vectors. In detail, *M/k*-1 pattern vectors were assigned to a training set to train the SVM. The withheld pattern vectors were assigned to a testing set and used to assess the performance of the trained SVM (% decoding accuracy). The training and testing procedure was repeated 100 times with random assignment of raw pattern vectors to averaged pattern vectors. For the case of reduced sensor data sets, this also involved resampling the sensors for each iteration to obtain an unbiased estimate of decoding accuracy. For each time point, we stored the classification result averaged across iterations in matrices of 92 × 92 size, indexed in rows and columns by the classified conditions. This decoding matrix is symmetric and has an undefined diagonal (no classification within condition).

#### 3.4.2 Time-resolved object category discrimination

We evaluated when MEG and EEG activation patterns allow discrimination of five different object categorizations at the super-ordinate (animate vs. inanimate, natural vs. artificial), ordinate (bodies vs. faces) and sub-ordinate category level (human vs. animal bodies and faces). For this, we partitioned the 92 × 92 decoding matrices into within- and between-category segments for the relevant categorization according to the pairs of conditions indexed by each matrix element. (Fig. 2A). The average of between minus within-category decoding accuracy values is a measure of clustering by category, indicating information about category membership over and above the discriminability of single images.

**Fig 2.**
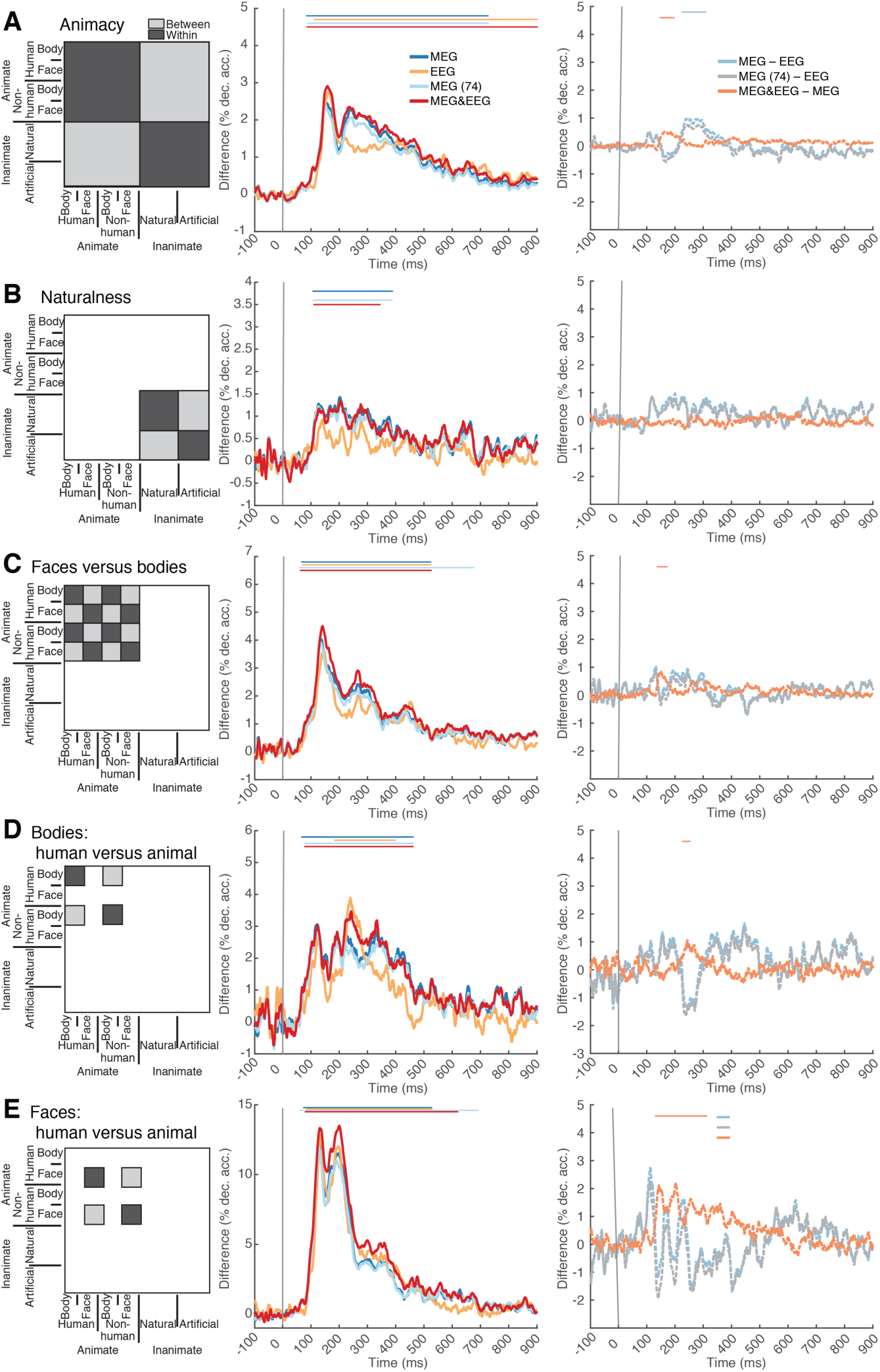
Time course of category membership from EEG and MEG data. Object category membership was assessed by representational clustering analysis for **A**) animacy, **B**) naturalness, **C**) faces vs. bodies, **D**) human versus nonhuman bodies and **E**) human versus nonhuman faces. For this, we partitioned the decoding matrix (left panels) in regions containing pairwise decoding accuracies within (dark gray) and between (light gray) the relevant categorical subdivisions (for peak latencies see Table 3). Right panels report the difference curves for results obtained from different samplings of MEG and EEG sensors (for peak latencies see Table 4). Lines above curves indicate significant time points (*n* = 15, cluster-defining threshold *P* < 0.05, corrected significance level *P* < 0.05 Bonferroni-corrected by number of plots for each subpanel, both two-sided). The gray vertical line indicates onset of image presentation. For equivalent results based on additional sensor samplings see Supplementary Fig. 3.

### 3.5 Common and unique aspects of visual representations in MEG and EEG data

To reveal the common versus unique aspects of visual representations captured by multivariate pattern analysis of MEG and EEG data, we used representational similarity analysis (RSA) (Fig. 3A). We interpret decoding accuracy as a dissimilarity measure (Cichy et al., 2014, 2016c, 2016a): the higher the decoding accuracy, the more dissimilar the activation patterns are for the classified conditions. Thus, MEG and EEG decoding matrices can be interpreted as representational dissimilarity matrices (RDMs) allowing a direct comparison between the two modalities. The basic idea is that if EEG and MEG measure similar signals, two objects that evoke similar patterns in EEG should evoke similar patterns in MEG, too.

**Fig. 3.**
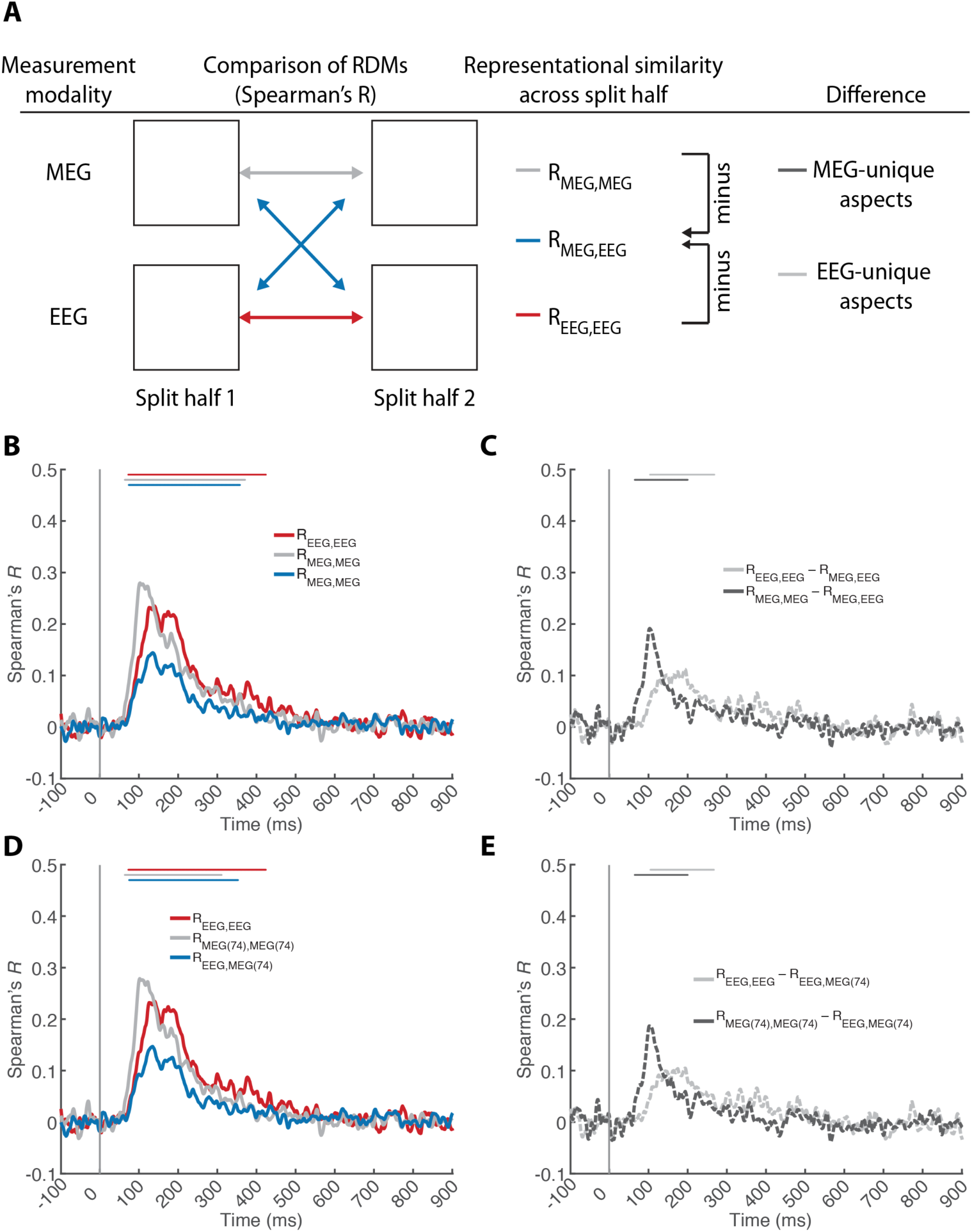
Time course of common and unique aspects of visual representations as resolved with MEG and EEG. **A**) Procedure. We split the MEG and EEG data in half (even and odd trials) to conduct two independent multivariate pattern classification analyses, yielding split-half specific RDMs. We then calculated representational similarity (Spearman’s *R*) across splits for the same measurement modality (MEG and EEG color-coded gray and red) and across modalities (color-coded blue). Comparing RDMs within imaging modalities resulted in a reliability estimate that includes both common and unique aspects of visual representations. Comparing RDMs across imaging modalities revealed the common aspects of visual representations. Thus the difference within-modality minus across-modality indicated the aspects of visual representations unique to each measurement modality (color-coded dark gray striped for MEG, and light gray striped for EEG. Within and across technique similarities are reported for **B,C**) EEG and MEG and **D,E**) EEG and MEG (74). Lines above curves indicate significant time points (*n* = 15, cluster-defining threshold *P* < 0.05, corrected significance level *P* < 0.05 Bonferroni-corrected by number of plots for each subpanel, both two-sided). The gray vertical line indicates image onset. For equivalent results based on additional sensor samplings see Supplementary Fig. 4.

A valid comparison of RDMs requires they are constructed from independent data (Henriksson et al., 2015). Otherwise, trial-by-trial signal fluctuations unrelated to experimental conditions, such as cognitive states (attention, vigilance) or external noise (movement, electromagnetic noise) will inflate, distort, and bias the similarity between EEG and MEG. For an independent construction of MEG and EEG RDMs we split the data in half by assigning even and odd trials to different sets. We then compared (Spearman’s *R*) the RDMs from split half 1 vs. split half 2 both within and across MEG and EEG measurement modalities using RSA (Fig 3A). Comparing RDMs across imaging modalities (MEG vs. EEG) revealed only the common aspects of visual representations. Comparing RDMs within imaging modalities (MEG vs. MEG and EEG vs. EEG; across data splits) resulted in a reliability estimate that includes both common and unique aspects. The difference of within-modality minus across-modality similarities thus revealed the unique aspects of visual representations measured with either MEG or EEG. For this analysis, the time-resolved classification was performed similarly to single image classification described above, but the sub-averaged pattern vectors were constructed by averaging *k* = 3 pattern vectors given the reduced number of trials.

### 3.6 fMRI stimulation protocol, acquisition, preprocessing and processing

We reanalyzed an existing fMRI data set reported in Cichy et al. (2014). Here we summarize the key points in fMRI data acquisition, preprocessing and processing for RSA-based fusion between fMRI and MEG/EEG data.

#### 3.6.1 Experimental paradigm

15 participants viewed the same 92 image set while fMRI data was recorded. Each participant completed two sessions on two separate days, where each session consisted of 10-14 runs of 384 s duration each. During each run every image was presented once, and image order was randomized. On each trial the image was shown for 500ms. The inter trial interval was 3 s. 25% of all trials were null trials during which only a gray background was presented, and the fixation cross turned darker for 100ms. Participants were instructed to report the change in fixation cross luminance with a button press.

#### 3.6.2 fMRI acquisition

We acquired MRI data on a 3T Trio scanner (Siemens, Erlangen, Germany) with a 32-channel head coil. Structural images were acquired in each session using a standard T1-weighted sequence (192 sagittal slices, FOV = 256 mm^2^, TR = 1,900 ms, TE = 2.52 ms, flip angle = 9°). Functional data were acquired with a gradient-echo EPI sequence (192 volumes, TR = 2,000 ms, TE = 31 ms, flip angle = 80°, FOV read = 192 mm, FOV phase = 100%, ascending acquisition, gap = 10%, resolution = 2 mm isotropic, slices = 25). The acquisition volume was partial and covered the ventral visual pathway.

#### 3.6.3 fMRI activation estimation

SPM8 (http://www.fil.ion.ucl.ac.uk/spm/) was used to process fMRI data. For each participant, we realigned fMRI data and co-registered it to the T1 structural scan acquired in the first MRI session. This formed the basis for the subsequent region-of-interest analysis. For the searchlight analysis (see below), fMRI data was additionally normalized to an MNI template. The subsequent processing was equivalent for both unnormalized and normalized data. To estimate the fMRI response to the 92 image conditions we used a general linear model (GLM). Onsets of image presentation entered the GLM as regressors and were convolved with the standard hemodynamic response function. Additional nuisance regressors were movement parameters and two regressors modelling session (1 for each volume of a session, 0 otherwise). Condition-specific GLM parameters (beta-values) were converted into *t*-values by contrasting each condition estimate against the implicitly modeled baseline. In addition, we modelled the overall effect of visual stimulation as a separate *t*-contrast of parameter estimates for all 92 conditions against baseline.

#### 3.6.4 fMRI region-of-interest definition

We assessed two regions-of-interest (ROIs): primary visual area V1 and inferior temporal cortex (IT). V1 was defined separately for each participant based on an anatomical eccentricity template, and contained all voxels assigned to the central 3 degrees of visual angle (Benson et al., 2012). IT was defined based on a mask consisting of bilateral fusiform and inferior temporal cortex (WFU PickAtlas, IBASPM116 Atlas (Maldjian et al., 2003)). To match V1 and IT ROIs in average size, we chose the 361 most activated voxels in the *t*-contrast of all image conditions vs. baseline.

#### 3.6.5 Region-of-interest-based fMRI representational similarity analysis

We constructed fMRI RDMs for each participant independently using a correlation-based dissimilarity measure. For each ROI we extracted and concatenated the fMRI voxel activation values for each image condition. We then calculated all pair-wise correlation coefficients (Pearson’s *R*) between the pattern vectors for each pair of image conditions and stored the result in a 92 × 92 symmetric matrix indexed in rows and columns by the compared conditions. We transformed the correlation similarity measure into a dissimilarity measure by subtracting the correlations coefficients from 1 (i.e., 1 – *R*). For further analyses, we averaged the resulting dissimilarity measures across sessions resulting in one RDM for each subject and ROI.

### 3.7 Spatial localization of MEG and EEG visual representations using fMRI-MEG/EEG fusion

To identify the spatial sources of the temporal dynamics observed in MEG and EEG, and to compare them to each other, we used a RSA-based MEG-fMRI fusion approach (Cichy et al., 2014, 2016c). The basic idea is that if locations resolved in fMRI and time points resolved in MEG/EEG correspond to each other, their corresponding RDMs should be similar.

#### 3.7.1 Region-of-interest-based fMRI-MEG/EEG fusion

For each ROI and subject we calculated the similarity (Spearman’s *R*) between the subject-specific fMRI RDM and the subject-averaged MEG or EEG RDM for each time point, yielding time courses (*n* = 15) of MEG-fMRI or EEG-fMRI representational similarity for each ROI and subject (Fig. 4A).

#### 3.7.2 Spatially unbiased searchlight fMRI-MEG/EEG fusion

For spatially unbiased fusion of fMRI with MEG and EEG beyond the ROI-based approach, we used a searchlight approach as introduced in Cichy et al. (2016) (Fig. 5a). We conducted the searchlight analysis separately for each fMRI subject (*n* = 15) and time point from -100 to +500 ms in 5 ms steps. For each voxel *v*, we extracted condition-specific *t*-value patterns in a sphere centered at *v* with a radius of 4 voxels (searchlight at *v*) and arranged them into pattern vectors. We calculated the pairwise dissimilarity between pattern vectors by 1 minus Pearson’s *R* for each pair of conditions, resulting in a fMRI RDM. We then calculated the similarity (Spearman’s *R*) between the searchlight-specific fMRI RDM and the subject-averaged MEG or EEG RDMs. Repeating this analysis for every voxel in the brain, we obtained a 3D map of representational similarities between fMRI and MEG or EEG at each time point. Repeating the same approach for all time points, we obtained a series of 3D maps revealing the spatio-temporal activation of the human brain during object perception as captured with MEG and EEG respectively.

### 3.8 Statistical testing

We conducted non-parametric random effects statistics throughout. We used permutation tests for cluster-mass inference, and bootstrap tests to determine confidence intervals of peak latencies (Nichols and Holmes, 2002; Pantazis et al., 2005; Maris and Oostenveld, 2007).

For the statistical assessment of the classification analysis, the MEG-EEG comparison by RSA, and the ROI-based fMRI-MEG/EEG fusion analysis we randomly shuffled the sign of the data points (10,000 permutation samples) for each subject to determine significant effects at a threshold of *P* < 0.05, two sided. To correct for multiple comparisons across voxels (fMRI) or time points (MEG/EEG), we used cluster-mass inference (i.e. number of significant elements weighed by the value of those elements) with a cluster extent threshold of *P* < 0.05). In addition, for multiple tests of the same hypothesis (as reported by a figure subpanel) we further Bonferroni corrected the cluster extent threshold.

The statistical assessment of the fMRI-MEG/EEG searchlight fusion analysis was as follows. To determine a cluster-defining threshold, we averaged the subject-specific fusion results (4-dimensional, i.e., 3 spatial × 1 temporal dimension) across subjects, and aggregated voxel values across space and time points from -100 to 0 ms to form an empirical baseline voxel distribution. When comparing representational similarity between fMRI and MEG/EEG, we determined the right-sided 99.99% threshold of the distribution, constituting a baseline-based cluster defining threshold at *P* < 0.001, one-sided. For comparison of results for different sensor samplings, we used an equivalent two-sided test procedure (*P* < 0.001, two-sided).

To obtain a permutation distribution of maximal cluster mass, we randomly shuffled the sign of subject-specific data (1,000 permutation samples). For each sample, we averaged data across subjects, and determined 4-dimensional mass of clusters (i.e. number of significant spatially and temporally connected elements weighed by their absolute value) exceeding the right-sided cluster threshold. We then determined the maximal cluster size. This yielded a distribution of maximal cluster sizes under the null hypothesis. We report clusters as significant if they were larger than the 95% threshold of the maximal cluster size distribution, corresponding to a *P* = 0.05 one-sided threshold. For two-sided tests, two distributions of maximal cluster size under the null were created, and clusters are reported as significant if they passed the 97.5% threshold, corresponding to a *P* = 0.05 two-sided threshold.

## 4 Results

### 4.1 Commonalities and differences in the time courses of single image classification from MEG and EEG data

We first investigated whether MEG/EEG signals allow for time-resolved discrimination of individual object images. For every time point, we averaged across all elements of the decoding matrices, yielding a time course of grand average decoding accuracy across all experimental conditions (Fig. 1C). We observed significant effects for all four main sensor samplings of MEG/EEG sensors (for peak latencies see Table 1; for references to Supplementary Tables and Figures reporting results based on supplementary sensor samplings please confer the figure and legend captions here and throughout the manuscript). This demonstrates that in principle both MEG and EEG signals lend themselves to the same kind of multivariate analysis, and reproduced the MEG-based results of Cichy et al. (2014).

**Table 1.** Peak latency of single image classification time courses for several samplings of MEG and EEG sensor data **A**), and differences therein **B**). Numbers in brackets indicate 95% confidence intervals. For equivalent results based on additional sensor samplings see Supplementary Table 1.

| Sensor Sampling | Peak latency (ms) |
| --- | --- |
| **A) Decoding results** |  |
| MEG | 112 (109 – 124) |
| EEG | 181 (131 – 195) |
| MEG (74) | 114 (109 – 125) |
| MEG&EEG | 114 (10 – 134) |
| **B) Differences in decoding results** |  |
| MEG – EEG | 107 (99 – 115) |
| MEG (74) – EEG | 107 (96 – 115) |
| MEG&EEG – MEG | 197 (177 – 201) |

We observed several differences in the EEG- and MEG-based time courses. First, classification accuracy for MEG was consistently higher than for EEG for most post-stimulus period. To quantify this effect, we subtracted the EEG from the MEG time course (Fig. 1D). Note that the higher number of sensors in the MEG analysis did not trivially explain this difference, as the reduced MEG (74) sensor data set yielded equivalent results (Fig. 1C,D; for details see Table 1). A second aspect in which MEG and EEG differed was peak latency: MEG-based time courses peaked significantly earlier than the EEG-based time course (P < 0.001, for details see Table 2, also independent of sensor number).

**Table 2.**
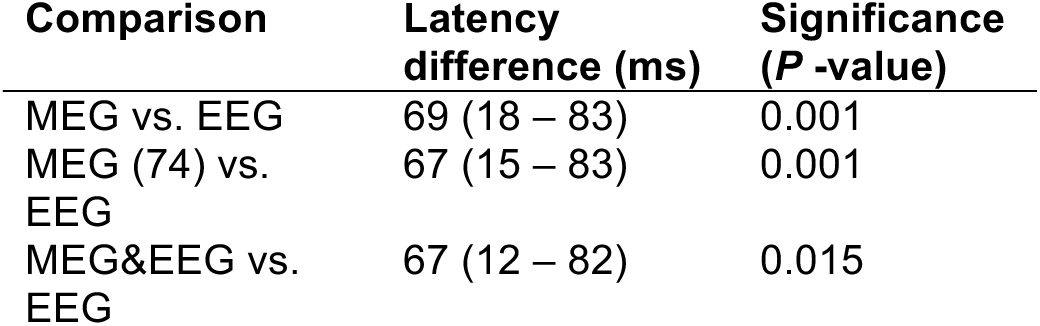
Comparison of peak latencies between single image classification time courses for different samplings of MEG and EEG sensor data. Significance was determined by bootstrapping the participant pool (*n* = 15, 10,000 bootstraps). Numbers in brackets indicate 95% confidence intervals. For equivalent results based on additional sensor samplings see Supplementary Table 2.

| Comparison | Latency difference (ms) | Significance (P -value) |
| --- | --- | --- |
| MEG vs. EEG | 69 (18 – 83) | 0.001 |
| MEG (74) vs. EEG | 67 (15 – 83) | 0.001 |
| MEG&EEG vs. EEG | 67 (12 – 82) | 0.015 |

In combination, the differences in grand average decoding and peak latency suggest that MEG and EEG may reflect partially different aspects of emerging visual representations. One prediction of this hypothesis is that combining MEG and EEG before multivariate pattern classification should yield higher decoding accuracy than MEG alone. We found this to be the case: the grand average decoding accuracy time course for combined MEG&EEG data was significantly higher than for MEG alone (Fig. 1 C,D).

In sum, we found that both MEG and EEG signals carry information at the level of single object images, but with differing temporal evolution suggesting sensitivity to partly different aspects of visual representations.

### 4.2 Time courses of visual category membership resolved with MEG and EEG are similar

Given the MEG and EEG qualitative and quantitative differences in decoding single images, we investigated whether MEG and EEG also differ in revealing information about object category processing at different levels of categorical abstraction. Following the same approach as in Cichy et al (2014), we partitioned the decoding accuracy matrix into two subdivisions (Fig. 2A-E, left panel): images belonging to the same (light gray) and to different (dark gray) subdivisions with respect to a particular categorization. The comparison of within and between average subdivision decoding accuracies serves as a measure of clustering by category membership. This is a measure of the explicitness of a representation, in the sense that category membership could be read out from it in linear fashion (DiCarlo and Cox, 2007).

We conducted this analysis for five different categorical subdivisions: at the super-ordinate category level for animacy (Fig. 2A) and naturalness (Fig. 2B), at the ordinate category level for faces vs. bodies (Fig. 2C) and at the sub-ordinate category level for human bodies vs. non-human bodies (Fig. 2D) and human faces vs. non-human faces (Fig. 2E). We found significant signals for category membership for all five subdivisions in all four samplings of MEG and EEG sensors (Fig 2A-E, middle panel, except naturalness in EEG (for details see Table 3). This result reinforces the point that multivariate pattern classification is similarly powerful when applied to EEG as when applied to MEG.

**Table 3.** Peak latency of category membership time courses for **A**) animacy, **B**) naturalness, **C**) face vs. body, **D**) human vs. animal face, and **E**) human vs. animal body. Numbers in brackets indicate 95% confidence intervals. For equivalent results based on additional sensor samplings see Supplementary Table 3.

| Sensor Sampling | Peak latency<br>(ms) |
| --- | --- |
| **A) animacy** |  |
| EEG | 159 (145 – 174) |
| MEG | 151 (146 – 292) |
| MEG&EEG | 157 (148 – 246) |
| MEG (74) | 152 (146 – 290) |
| **B) body vs. face** |  |
| EEG | 146 (127 – 171) |
| MEG | 132 (129 – 161) |
| MEG&EEG | 140 (132 – 162) |
| MEG (74) | 134 (128 – 159) |
| **C) human vs. animal body** |  |
| EEG | 239 (127 – 270) |
| MEG | 121 (114 – 359) |
| MEG&EEG | 241 (121 – 333) |
| MEG (74) | 123 (113 – 366) |
| **D) human vs. animal face** |  |
| EEG | 133 (128 – 202) |
| MEG | 128 (125 – 211) |
| MEG&EEG | 200 (126 – 210) |
| MEG (74) | 128 (124 – 212) |
| **E) natural vs. artificial** |  |
| EEG | 248 (133 – 585) |
| MEG | 202 (116 – 307) |
| MEG&EEG | 203 (125 – 638) |
| MEG (74) | 204 (115 – 303) |

Analogous to the investigation of the grand average time courses above, we investigated differences between MEG and EEG based results in decoding accuracy, differences in peak latency, and whether combining MEG&EEG signals yielded higher decoding accuracy than MEG alone. Concerning the difference between category-specific curves derived from MEG and EEG data (Fig. 2, right panels), we found only minor and transient statistical differences (for details see Table 4). Comparing peak latency differences, we found no significant effects (all *P* > 0.12). Finally, the comparison of the results based on sampling MEG&EEG vs. MEG revealed a difference in all cases, except for naturalness (Fig. 2A-E).

**Table 4.**
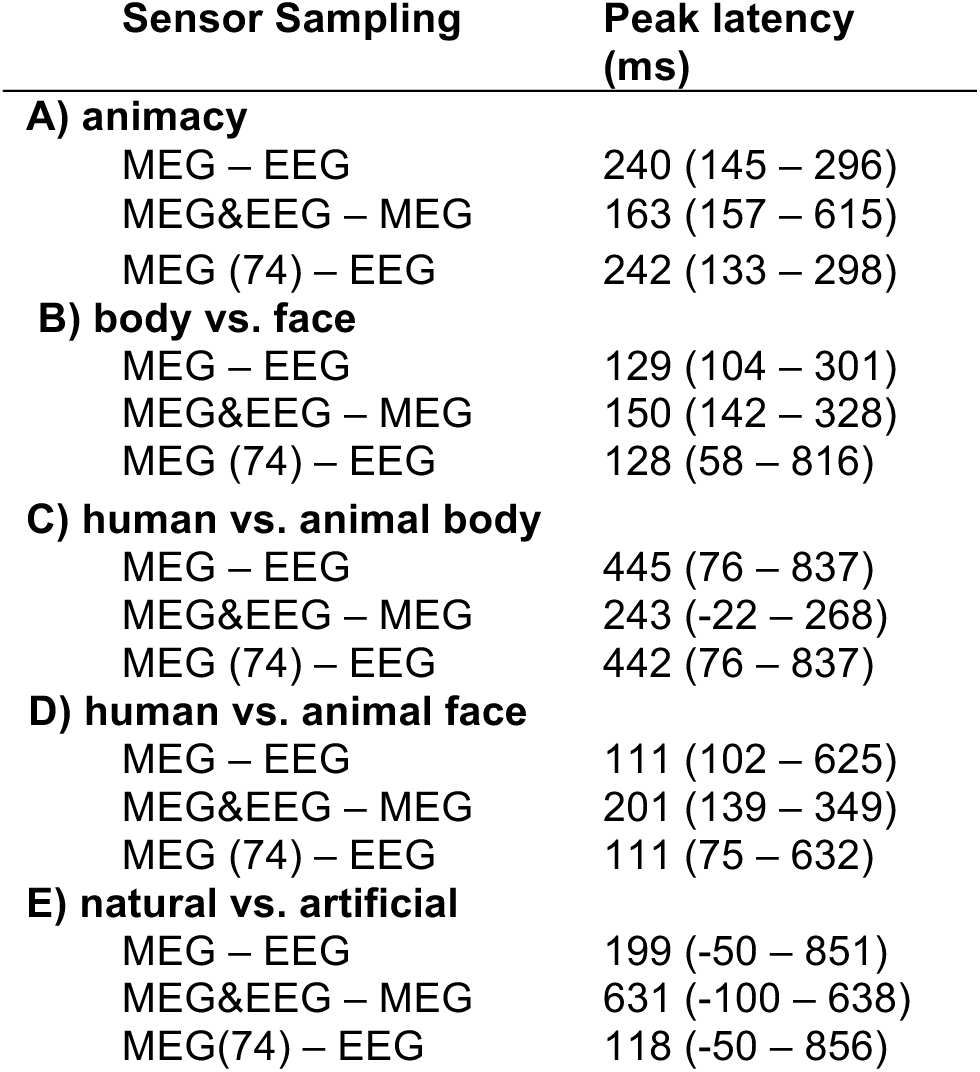
Peak latency of differences of category membership time courses for **A**) animacy, **B**) naturalness, **C**) face vs. body, **D**) human vs. animal face) and **E**) human vs. animal body. For equivalent results based on additional sensor samplings see Supplementary Table 4.

Together, these results show that for category-specific signals, MEG and EEG resolve visual representations with similar time courses, and further suggest that MEG and EEG may be partially sensitive to different aspects of visual representations.

### 4.3 Comparison of MEG and EEG data by representational similarity analysis revealed both common and unique aspects of neural representations

Grand average single image decoding accuracy and category-specific signals are summary statistics that only partially reflect the rich multivariate information in MEG and EEG data. How do MEG and EEG compare if the entire structure of representational space captured by the decoding matrices is considered?

To investigate, we used representational similarity analysis (RSA) (Kriegeskorte, 2008; Kriegeskorte and Kievit, 2013b) on the full decoding matrix. The idea is that decoding accuracy can be seen as a dissimilarity measure: condition pairs that have similar representations yield low decoding accuracy, and condition pairs that have dissimilar representations yield high decoding accuracies (Cichy et al., 2014, 2016a). The decoding matrices for MEG and EEG can be thus interpreted as representational dissimilarity matrices (RDMs), summarizing similarity relations between sensor activation patterns related to experimental conditions. MEG and EEG decoding matrices can then be compared directly for similarity. Importantly, to yield an unbiased measure of similarity, RDMs must be based on brain data recorded independently, i.e. for different trials (Henriksson et al., 2015). We thus split the MEG and EEG data in half (even versus odd trials), and conducted multivariate pattern classification based on each split half data set, equivalently to the analysis of the full data as explicated above. All RDM comparisons (Spearman’s *R*) were then conducted across split halves (Fig. 3A).

Comparing RDMs across imaging modalities (R_MEG,EEG_) revealed the common aspects of visual representations (Fig. 3B, blue line, for peak latencies see Table 5A). We found a positive and significant representational similarity time course, indicating aspects of visual representations resolved by both modalities. Comparing RDMs within imaging modalities (R_MEG,MEG_ and R_EEG,EEG_) resulted in a reliability estimate that includes both common and unique aspects of visual representations (Fig 3B, gray and red line respectively). These were also significant, and notably, higher than the across-modality representational similarities, indicating that MEG and EEG resolve partly unique aspects of visual representations. The difference of within-modality minus across-modality similarity curves, a measure that quantifies the unique information in each modality, statistically ascertained this result (Fig. 3C, for details see Table 5A).

**Table 5.**
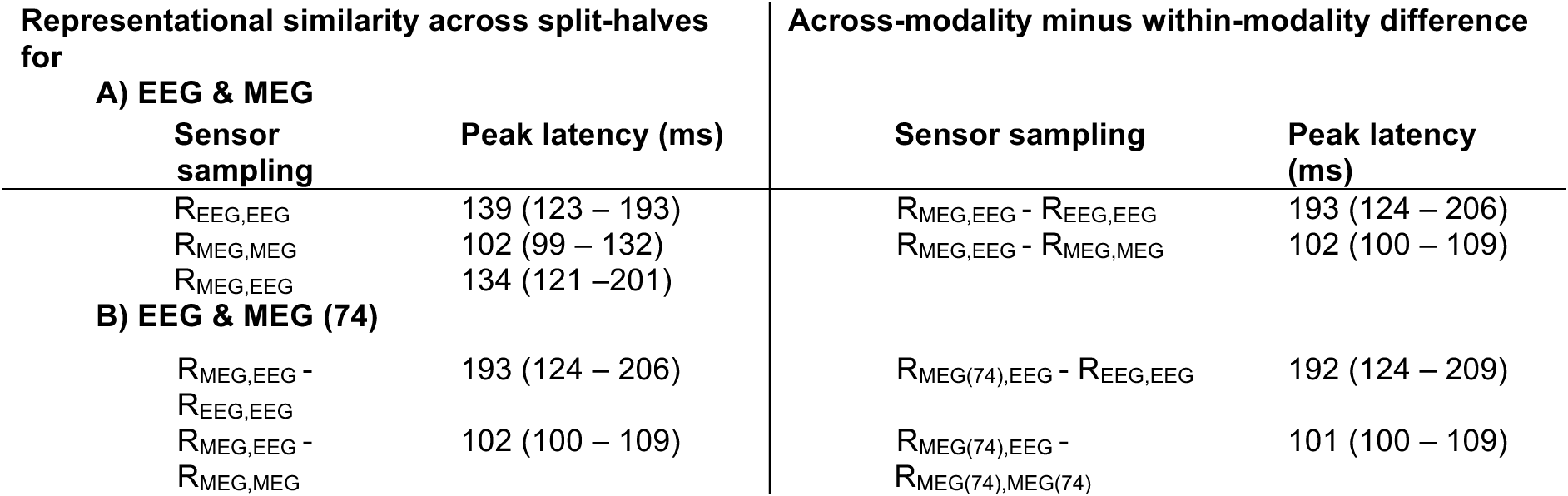
Peak latency of representational similarity time courses within and across measurement modalities (EEG and MEG) (left half) and differences of across-modality minus within-modality similarities (right half) for **A**) full sensor set, and **B**) MEG (74). Numbers in brackets indicate 95% confidence intervals. For equivalent results based on additional sensor samplings see Supplementary Table 5.

| Representational similarity across split-halves for |  | Across-modality minus within-modality difference |  |
| --- | --- | --- | --- |
| A) EEG & MEG |  |  |  |
| Sensor sampling | Peak latency (ms) | Sensor sampling | Peak latency (ms) |
| R<sub>EEG,EEG</sub> | 139 (123 – 193) | R<sub>MEG,EEG</sub> - R<sub>EEG,EEG</sub> | 193 (124 – 206) |
| R<sub>MEG,MEG</sub> | 102 (99 – 132) | R<sub>MEG,EEG</sub> - R<sub>MEG,MEG</sub> | 102 (100 – 109) |
| R<sub>MEG,EEG</sub> | 134 (121 – 201) |  |  |
| B) EEG & MEG (74) |  |  |  |
| R<sub>MEG,EEG</sub> - R<sub>EEG,EEG</sub> | 193 (124 – 206) | R<sub>MEG(74),EEG</sub> - R<sub>EEG,EEG</sub> | 192 (124 – 209) |
| R<sub>MEG,EEG</sub> - R<sub>MEG,MEG</sub> | 102 (100 – 109) | R<sub>MEG(74),EEG</sub> - R<sub>MEG(74),MEG(74)</sub> | 101 (100 – 109) |

The time course of MEG- and EEG-unique signals was different: the peak latency was significantly earlier for MEG than for EEG (Δ = 91ms; *P* = 0.0003). Importantly, this result was not dependent on sensor number differences, as equivalent results were obtained when equating the number of MEG and EEG sensors (Δ = 91ms; P = 0.0001; Fig. 3D,E, Table 5B).

Together, these results demonstrate that MEG and EEG resolve partially common, and partially unique aspects of visual representations in the brain, with differentiable temporal dynamics for unique components.

### 4.4 Fusion with fMRI revealed the locus of unique and common aspects of visual representations resolved with MEG and EEG

To investigate the cortical locus of the unique and common aspects of neural representations resolved by MEG and EEG as identified above, we used the fMRI-MEG/EEG fusion approach proposed in Cichy et al., 2014. By objectively evaluating MEG and EEG data against an independent data set of fMRI, fusion bypasses the inherent ambiguities of spatial localization methods relying on MEG/EEG alone.

We first investigated the source of EEG and MEG signals in the ventral visual stream per se. For this, we compared the representational similarity between fMRI-based RDMs for two cortical regions – early visual cortex (V1) and inferior temporal cortex (IT) – with the time-resolved RDMs for MEG and EEG respectively (Fig. 4A).

We found significant fMRI and MEG/EEG representational similarities in both V1 and IT for all the investigated sensor samplings of MEG and EEG (Fig. 4B,D, for details see Table 6). Consistent with the view of visual processing as a spatiotemporal cascade along the ventral visual stream, representational similarities between fMRI and MEG/EEG signals peaked earlier for V1 than for IT (for all sensor samplings, P < 0.01, Bonferroni-corrected for multiple comparisons). This reproduces previous results from MEG-fMRI fusion (Cichy et al., 2014), extends them to EEG, and reinforces the view of visual processing as a spatio-temporal cascade from posterior to anterior visual regions over time.

**Fig. 4.**
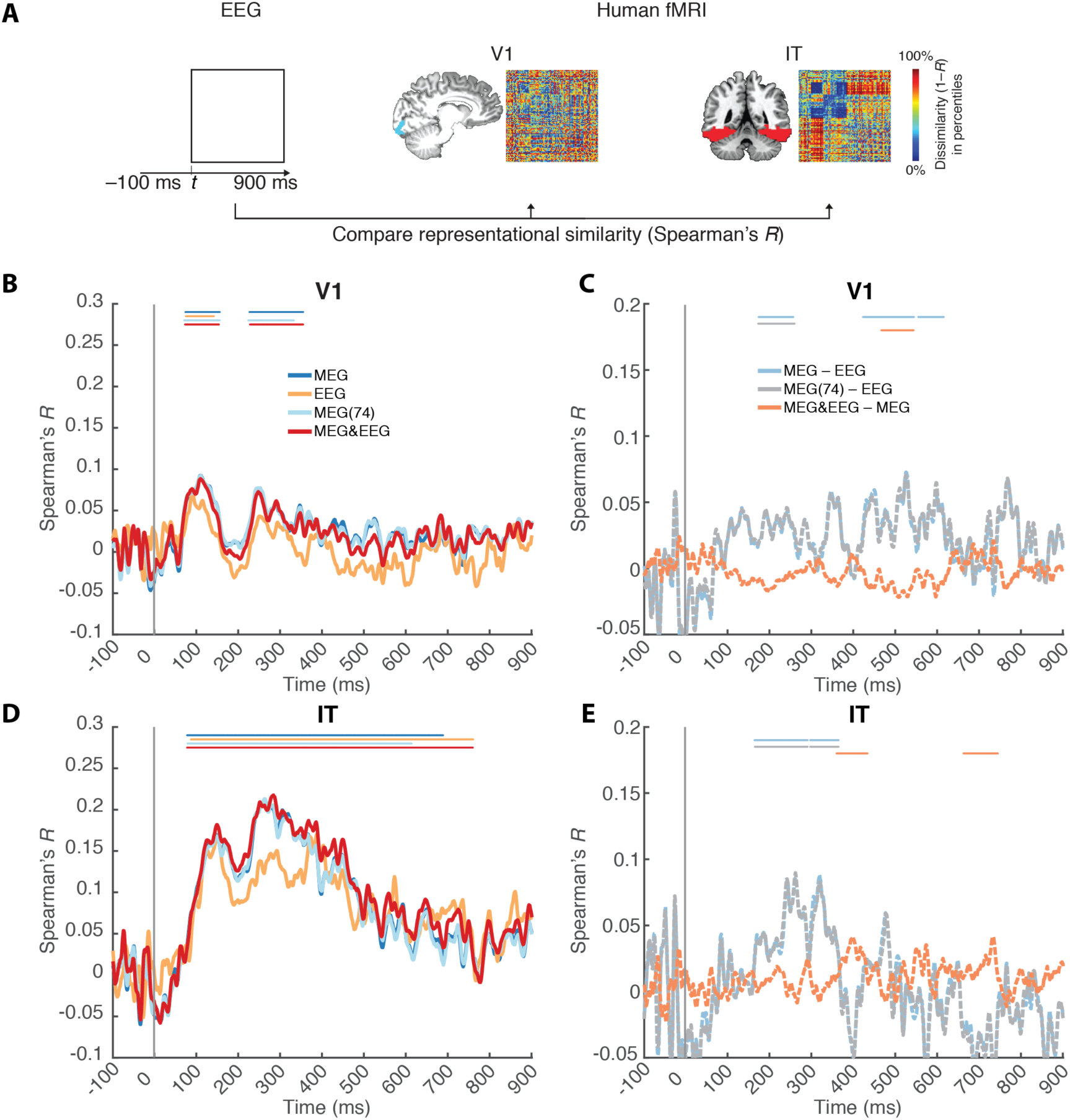
Relating fMRI to MEG/EEG signals using a ROI-based fusion approach. **A)** Procedure. For two regions-of-interest (V1, IT) we calculated representational dissimilarity matrices (fMRI RDMs) based on fMRI activation patterns to the same image set as in the MEG and EEG data. In detail, for every time point t, we calculated the similarity (Spearman’s *R*) between fMRI and MEG or EEG RDMs. This resulted in time courses of representational correspondence between MEG/EEG and fMRI in **B**) V1 and **D**) IT. MEG and EEG had similar time courses, with representational correspondence emerging earlier in time for V1 than for IT. **C,E)** Difference curves for results reported in B and D, revealing stronger signals for MEG than for EEG. Lines above curves indicate significant time points (*n* = 15, cluster-defining threshold *P* < 0.05, corrected significance level *P* < 0.05 Bonferroni-corrected by number of plots for each subpanel, both two-sided). The gray vertical line indicates onset of image presentation. For equivalent results based on additional sensor samplings see Supplementary Fig. 5.

**Table 6.** Peak latency for **A**) ROI-based fMRI-MEG/EEG fusion results for V1 and IT, and **B**) difference curves for different sensor samplings. Numbers in brackets indicate 95% confidence intervals. For equivalent results based on additional sensor samplings see Supplementary Table 6.

| Sensor Samplings | V1 | IT |
| --- | --- | --- |
| **A) fMRI fusion results with** |  |  |
| EEG | 89 (87 – 125) | 387 (145 – 388) |
| MEG | 109 (81 – 245) | 262 (250 – 306) |
| MEG&EEG | 109 (78 – 247) | 283 (250 – 286) |
| MEG(74) | 109 (78 – 246) | 262 (259 – 282) |
| **B) Differences in fMRI-MEG/EEG fusion results** |  |  |
| MEG – EEG | 525 (-25 – 770) | 318 (-50 – 321) |
| MEG(74) – EEG | 525 (-26 – 769) | 261 (-50 – 321) |
| MEG&EEG – MEG | -17 (-17 – 734) | 733 (-18 – 733) |

We next investigated whether the representational similarity between MEG/EEG and fMRI patterns in V1 and IT is due to unique or common aspects in MEG or EEG in three analyses. First, we compared peak latency differences for the different samplings of MEG and EEG data. We found no significant differences (all *P* > 0.14 bootstrap across participants, 10,000 iterations), suggesting that potential differences between MEG and EEG are not to be found in the shape of the time courses.

Second, we subtracted EEG- from MEG-based fusion results (Fig. 4C,D) to determine which modality bore closer similarity to the fMRI patterns. Overall, we found MEG-based fusion consistently stronger than EEG-based fusion. However, comparison of MEG&EEG versus MEG alone produced inconsistent results with opposite sign for V1 and IT (for details see Table 6). Thus, this analysis did not reveal consistent difference either.

Third, for a particularly strong and sensitive test, we used a partial correlation analysis to investigate the relation between fMRI and MEG/EEG when the effect of either modality is partialled out (Fig. 5A, example of fMRI-EEG fusion in V1 when partialling out the MEG RDM). Such analysis should reveal representational similarities specific to a modality, by controlling for the effects of the other modality. We found that for V1, partialling out MEG from EEG abolished the significant representational correspondence to fMRI, whereas partialling out EEG from MEG did not (for details see Table 7). This suggests that MEG is sensitive to unique sources in V1 as compared to EEG. For IT, we found stable and significant representational correspondence for both MEG and EEG with fMRI when the effect of either EEG or MEG was accounted for. This shows that MEG and EEG both resolve unique aspects of representations in IT.

Overall, these results demonstrate that both MEG and EEG are well suited for RSA-based fusion analysis with fMRI. While both MEG and EEG are sensitive to unique aspects of visual representations in IT, only MEG is sensitive to unique aspects of visual representations in V1.

**Fig. 5.**
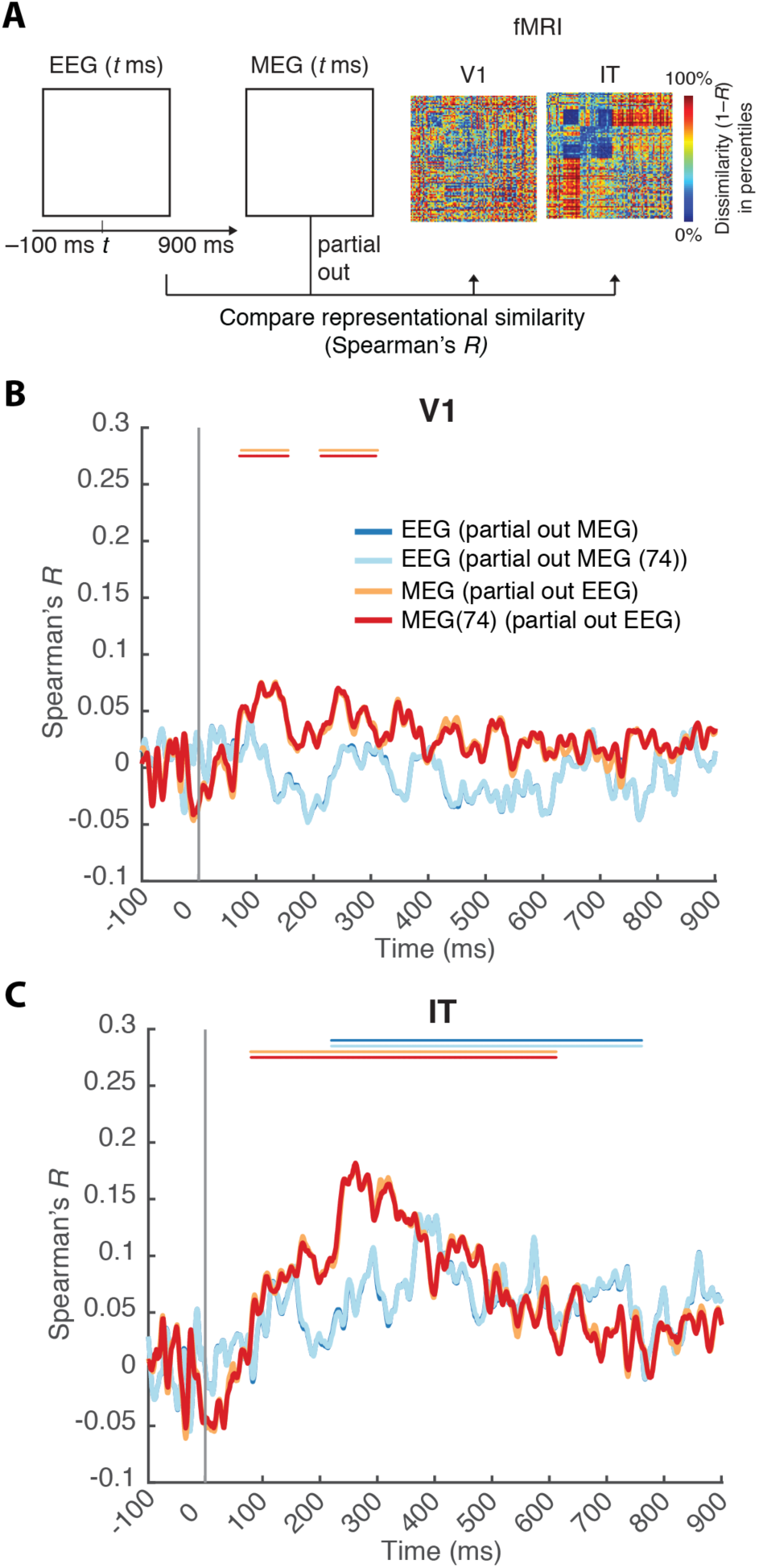
Relating fMRI to MEG/EEG signals using a ROI-based fusion approach and partial correlation analysis. **A**) Procedure. For every time point *t*, we calculated the similarity (Spearman’s *R*) between fMRI and EEG RDMs while partialling out the MEG RDMs. Equivalent Analyses were conducted for different MEG/EEG sensor samplings. Results of the partial correlation analysis are reported for **B**) V1 and **C**) IT. Lines above curves indicate significant time points (*n* = 15, cluster-defining threshold *P* < 0.05, corrected significance level *P* < 0.05 Bonferroni-corrected by number of plots for each subpanel, both two-sided). The gray vertical line indicates image onset. For equivalent results based on additional sensor samplings see Supplementary Fig. 6.

**Table 7.** Peak latency of the ROI-based fMRI-MEG/EEG fusion results after partialling out the effects of the other modality. For equivalent results based on additional sensor samplings see Supplementary Table 8.

| Fusion analysis | V1 | IT |
| --- | --- | --- |
| EEG (partial out MEG) | - | 371 (371 – 572) |
| EEG (partial out MEG(74)) | - | 371 (371 – 572) |
| MEG (partial out EEG) | 133 (80 – 253) | 262 (242 – 318) |
| MEG(74) (partial out EEG) | 108 (78 – 251) | 261 (241 – 282) |

### 4.5 MEG and EEG equally resolved the spatiotemporal dynamics of the ventral pathway as revealed by searchlight-based fusion with fMRI

What are the sources of MEG/EEG activity beyond the two investigated ROIs V1 and IT? To create a spatially unbiased view of the spatiotemporal dynamics in the ventral pathway, we used a searchlight-based fusion analysis (Fig. 6A). In particular, we investigated whether the fusion of fMRI with MEG, introduced in Cichy et al. (2016), can be directly extended to EEG, and whether such approach can reveal MEG and EEG differences beyond V1 and IT.

**Fig. 6.**
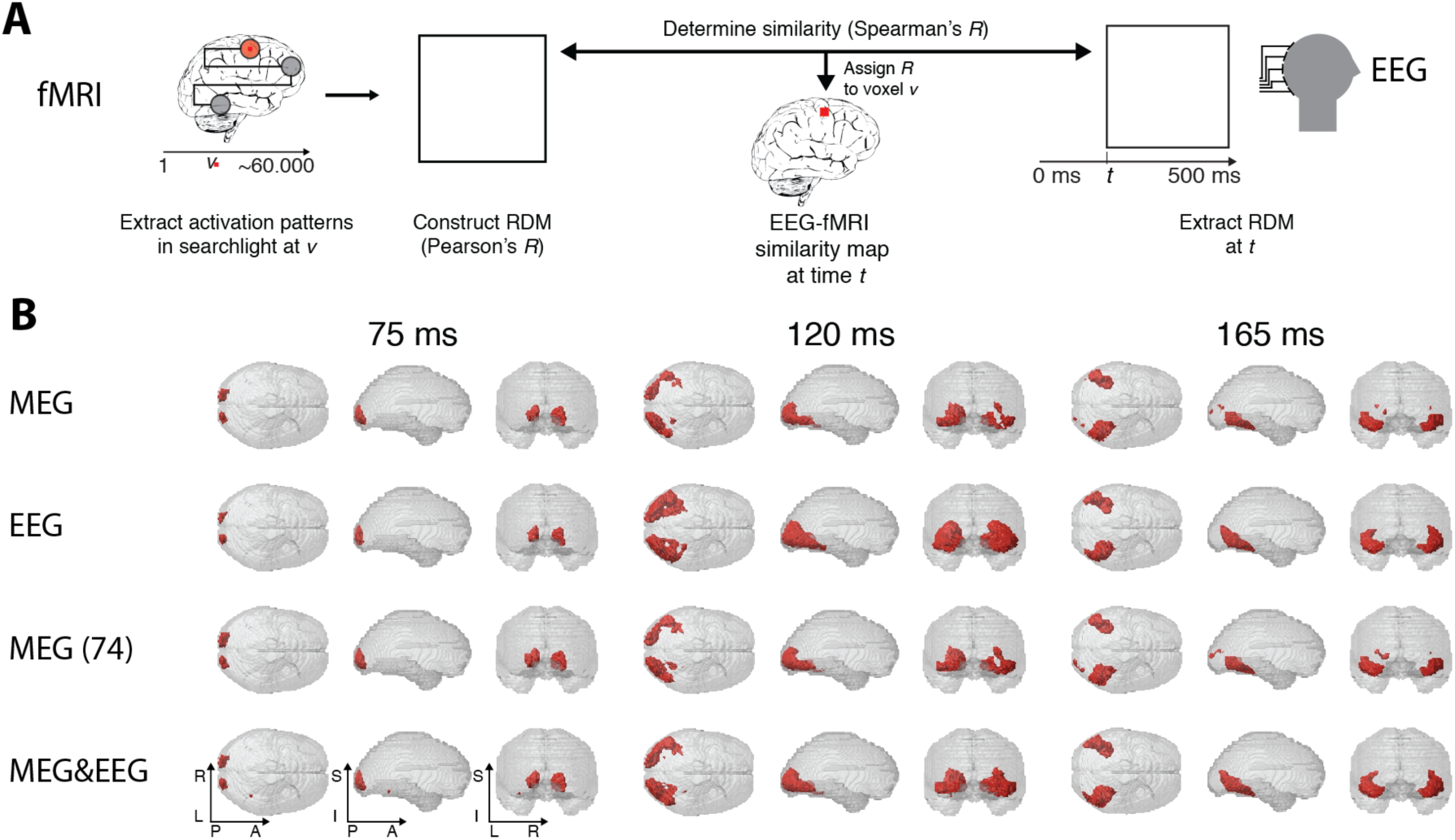
Spatially unbiased fusion analysis of fMRI with MEG and EEG. **A**) Procedure. Time-resolved MEG/EEG RDMs were compared (Spearman’s *R*) to space-resolved fMRI RDMs derived with searchlight analysis. In particular, for each voxel v in the brain we extracted fMRI activation patterns in the local vicinity of the voxel and calculated fMRI RDMs based on these patterns. Repeating for each voxel in the brain, this yielded a map of fMRI RDMs across the whole brain. Fusion of the time-resolved MEG/EEG RDMs with the space-resolved fMRI RDMs produced the spatiotemporal dynamics of representational correspondence. **B**) Snapshots at 75, 120 and 165ms for EEG, MEG, MEG(74) and MEG&EEG. All analyses revealed comparable spatio-temporal dynamics during object vision in the ventral visual stream: representational correspondence emerged first at the occipital pole, before extending rapidly anterior along the ventral visual pathway. Red voxels indicate statistical significance (*n* = 15, cluster-definition threshold *P* < 0.001, cluster threshold *P* < 0.05, both two-sided). A time-resolved movie is available (Supplementary movie 1). Inset axis indicate orientation of transparent brains (L/R= left/right; P/A = posterior/anterior; I/S = inferior/superior).

Both MEG and EEG-based fusion with fMRI data revealed a feed-forward cascade of representational similarity in the ventral visual stream (Fig. 6B): early representational relations were similar in the occipital pole, rapidly spreading along the ventral visual stream with comparable dynamics. This reproduced previous findings with MEG, and demonstrated the feasibility of the spatially unbiased searchlight-based fusion approach with EEG. Equivalent results were found for the reduced 74-sensor MEG data set, as well as combining MEG and EEG data prior to fusion (Fig. 6B).

We next compared the fusion results across the different MEG and EEG data sets. There were no significant effects for MEG versus EEG, irrespective of using the complete or the reduced 74-sensor MEG array (cluster definition threshold *P* < 0.001 cluster threshold *P* < 0.05, two sided). Similarly, comparison of MEG versus MEG&EEG fusion-based analysis did not yield significant results.

Together, the results demonstrate that both MEG and EEG are well suited for RSA-based fusion with fMRI data to reveal cortical information flow, but did not reveal further sources of sensitivity of MEG/EEG to unique aspects of visual representations.

## 5 Discussion

### 5.1 Summary

To investigate how relative sampling differences of neural activity inherent to MEG and EEG impact multivariate pattern analysis, we compared concurrently acquired MEG and EEG data. We found that all analyses yielding significant results in one measurement modality yielded significant results in the other modality, too. Comparison of MEG and EEG by classification-based time courses, as well as directly by representational similarity analysis yielded evidence for sensitivity to both common as well as unique aspects of neural representations. Fusion of MEG and EEG with fMRI localized the unique aspects: both modalities captured unique aspects of representations in high-level visual cortex, and MEG also in early visual cortex.

### 5.2 Both EEG and MEG are well suited for multivariate analyses methods to reveal human cortical dynamics

A recently and rapidly emerging range of research efforts have applied multivariate analyses to both EEG and MEG, demonstrating that both modalities resolve information about a diverse array of sensory and cognitive processes. These include discrimination of visual stimuli during passive viewing (Carlson et al., 2013; Cichy et al., 2014; Clarke et al., 2014; Kaneshiro et al., 2015), associative retrieval and learning (Kurth-Nelson et al., 2016), the emergence of invariance to changes in viewing conditions (Isik et al., 2014; Kietzmann et al., 2016), the temporal stability of neural signals (King and Dehaene, 2014), the maintenance of neural representations in dual tasks (Marti et al., 2015), and the comparison of neural representations with computational models (Groen et al., 2013; Cichy et al., 2016a), and across imaging modalities (MEG-fMRI) (Cichy et al., 2016c) and species (human-monkey) (Cichy et al., 2014).

While the successful application of multivariate methods to EEG and MEG in isolation demonstrates the potential of both modalities, it leaves open how the two relate, i.e. whether one modality is better suited for a particular analysis, and whether the results obtained from MEG and EEG measurements are comparable. Here we have shown in direct comparison of MEG and EEG for a large range of analyses, ranging from single image classification to fusion with fMRI data based on representational similarity analysis, they yield largely convergent results.

Our results have several practical implications for the application of multivariate analysis methods to MEG and MEG. As the availability and number of EEG devices vastly surpasses the number of MEG facilities, we hope that the observed comparability in results greatly increases the reach of this methodology. Moreover, most effects were robustly detectable when even only 32 EEG electrodes were sampled (Supplementary Figures 2,3,5,6). This shows that multivariate pattern analysis methods are suitable tools even when only a low numbers of sensors are recorded, e.g. in clinical settings. Further, in most analyses the effect size was smaller for EEG than MEG. This suggests that if MEG and EEG are equally accessible and time is a hard constraint, MEG might be preferred. If time is not a constraint, long EEG recording times may possibly offset any MEG advantage. It is also conceivable that modern high-density EEG electrode arrays, reaching a few hundred sensors, offer considerably improved results than our 74 passive electrode setup.

We hope that our results will motivate further researchers to widely use multivariate methods such as classification and RSA on both EEG and MEG data to shed further insight into the spatial and temporal neural dynamics underlying human cognition.

### 5.3 Multivariate analysis of MEG and EEG reveals both common and unique aspects of neural representations

Previous studies have yielded qualitatively comparable results for several kinds of multivariate analyses in MEG and EEG independently (Carlson et al., 2012; Cichy et al., 2014; Kaneshiro et al., 2015), thus suggesting sensitivity to common aspects of neural representations for MEG and EEG. However, for quantitative comparison and isolation of common and unique components it is important to establish correspondence in the same subjects recorded under the same experimental conditions. Otherwise, factors that differ across subjects or recording times may bias the comparison, including internal factors such as attention, vigilance, movement, cortical folding patterns, as well as differences in external conditions, such as noise and visual stimulation conditions.

Using multivariate analyses methods on concurrently recorded MEG and EEG data, we avoided those pitfalls, and presented corroborating evidence that MEG and EEG capture common and unique aspects of neural representations. Strongest evidence for both common and unique aspects was provided by direct comparison of MEG and EEG split-half data through RSA, which reveal within- and across-modality representational similarities millisecond by millisecond.

Further evidence specifically for unique aspects were differences in the time course of single image decoding and MEG/EEG-fMRI fusion, with later peaks for EEG than for MEG. Fusion of MEG/EEG with fMRI suggested that differential sensitivity to unique aspects of neural representations in early and late visual areas might explain this pattern. Only MEG revealed unique aspects in early visual areas, whereas both measurement modalities did so for later visual areas.

Last, the observation that effect sizes were larger when MEG and EEG data were combined, rather than used in separation, is consistent with the hypothesis that MEG and EEG are sensitive to unique aspects. However, an alternative possibility is that the gain in effect size was due to an increase of signal-to-noise ratio by combining measurements of common aspects with partially independent noise. Future in-depth quantitative evaluation and modelling efforts to equate noise levels across sensor sets are necessary to rule out this alternative explanation.

What are the reasons for the observed differences between MEG and EEG, and how do our findings relate to previous research? While MEG and EEG signals have the same underlying neural generators, there are well known and systematically explored differences in the literature (Cuffin and Cohen, 1979; Cohen and Cuffin, 1983; Hämäläinen et al., 1993). Prominently, radially-oriented sources are prominent in EEG but nearly silent in MEG, suggesting unique sensitivity of EEG to neural representations that are radially oriented. Additionally, EEG has higher sensitivity to deep sources than MEG, thus suggesting unique sensitivity to representations in cortical regions far away from the sensors. In contrast, volume currents measured by EEG are deflected and smeared by the inhomogeneity of resistance of the skull, scalp, and the different tissues of the human brain, potentially mixing signals from different sources more than MEG. Together, those differences make plausible the reasons for differential sensitivity of MEG and EEG in multivariate pattern classification, too.

In particular, a large body of theoretical, practical, and experimental investigations exploring the complementary nature of MEG and EEG data agrees with our observation that combining MEG and EEG increases effect size. Theoretical investigations predict the benefits of MEG/EEG data integration (Fokas, 2009). Practical and experimental investigations showed that combining MEG and EEG improves source reconstruction (Fuchs et al., 1998; Baillet et al., 1999; Pflieger et al., 2000; Liu et al., 2002; Yoshinaga et al., 2002; Liu et al., 2003; Babiloni et al., 2004; Huang et al., 2007; Sharon et al., 2007; Molins et al., 2008; Fokas, 2009; Henson et al., 2009).

Finally, our results suggest a potential future venue for the study of the complementarity of MEG and EEG responses. One pertinent prediction of the selective sensitivity of EEG to radial sources is that in a fusion-based comparison to fMRI, representations in cortical areas oriented radially should show stronger representational similarity to EEG than to MEG. Fusion-based analysis with custom designed fMRI RDMs selective of voxel patterns with preference to radial or tangential sources could improve localization and highlight the differential sensitivity of MEG and EEG signals to tangential and radial sources. Note, this is beyond the scope of this study, as fMRI was recorded in different subjects than EEG/MEG, making such and individualized analysis based on cortical folding patterns impossible.

## 6 Acknowledgements

We are grateful to Yu-Teng Chang and Yasaman Bagherzadeh for assistance in MEG and EEG data collection, and Martin Hebart and Matthias Guggenmos for comments on a previous version of this manuscript. This work was funded by the German Research Foundation (DFG, CI241/1-1) to R.M.C. and by the McGovern Institute Neurotechnology Program (to D.P.). MEG and EEG data were collected at the Athinoula A. Martinos Imaging Center at the McGovern Institute for Brain Research, MIT.

## References

Babiloni F, Babiloni C, Carducci F, Romani GL, Rossini PM, Angelone LM, Cincotti F (2004) Multimodal integration of EEG and MEG data: A simulation study with variable signal-to-noise ratio and number of sensors. Hum Brain Mapp 22:52–62.

Baillet S, Garnero L, Marin G, Hugonin JP (1999) Combined MEG and EEG source imaging by minimization of mutual information. IEEE Trans Biomed Eng 46:522–534.

Benson NC, Butt OH, Datta R, Radoeva PD, Brainard DH, Aguirre GK (2012) The Retinotopic Organization of Striate Cortex Is Well Predicted by Surface Topology. Curr Biol 22:2081–2085.

Carlson T, Alink A, Tovar D, Kriegeskorte N (2012) The evolving representation of objects in the human brain. J Vis 12:272–272.

Carlson T, Tovar DA, Alink A, Kriegeskorte N (2013) Representational dynamics of object vision: The first 1000 ms. J Vis 13:1–19.

Chang C, Lin C (2011) LIBSVM: a library for support vector machines. ACM Trans Intel Sys Tech, 2:27–27, 2011.

Cichy RM, Khosla A, Pantazis D, Oliva A (2016a) Dynamics of scene representations in the human brain revealed by magnetoencephalography and deep neural networks. Neuroimage doi: 10.1016/j.neuroimage.2016.03.063.

Cichy RM, Khosla A, Pantazis D, Torralba A, Oliva A (2016b) Comparison of deep neural networks to spatio-temporal cortical dynamics of human visual object recognition reveals hierarchical correspondence. Sci Reports doi:10.1038/srep27755

Cichy RM, Pantazis D, Oliva A (2014) Resolving human object recognition in space and time. Nat Neurosci 17:455–462.

Cichy RM, Pantazis D, Oliva A (2016c) Similarity-Based Fusion of MEG and fMRI Reveals Spatio-Temporal Dynamics in Human Cortex During Visual Object Recognition. Cereb Cortex:bhw135.

Clarke A, Devereux BJ, Randall B, Tyler LK (2014) Predicting the Time Course of Individual Objects with MEG. Cereb Cortex 25:3602–3612.

Cohen D, Cuffin BN (1983) Demonstration of useful differences between magnetoencephalogram and electroencephalogram. Electroencephalogr Clin Neurophysiol 56:38–51.

Cohen D, Hosaka H (1976) Part II magnetic field produced by a current dipole. J Electrocardiol 9:409–417.

Cuffin BN, Cohen D (1979) Comparison of the magnetoencephalogram and electroencephalogram. Electroencephalogr Clin Neurophysiol 47:132–146.

DiCarlo JJ, Cox DD (2007) Untangling invariant object recognition. Trends Cogn Sci 11:333–341.

Fokas AS (2009) Electro–magneto-encephalography for a three-shell model: distributed current in arbitrary, spherical and ellipsoidal geometries. J R Soc Interface 6:479–488.

Fuchs M, Wagner M, Wischmann H-A, Köhler T, Theißen A, Drenckhahn R, Buchner H (1998) Improving source reconstructions by combining bioelectric and biomagnetic data. Electroencephalogr Clin Neurophysiol 107:93–111.

Groen IIA, Ghebreab S, Prins H, Lamme VAF, Scholte HS (2013) From Image Statistics to Scene Gist: Evoked Neural Activity Reveals Transition from Low-Level Natural Image Structure to Scene Category. J Neurosci 33:18814–18824.

Hämäläinen M, Hari R, Ilmoniemi RJ, Knuutila J, Lounasmaa OV (1993) Magnetoencephalography—theory, instrumentation, and applications to noninvasive studies of the working human brain. Rev Mod Phys 65:413–497.

Henriksson L, Khaligh-Razavi S-M, Kay K, Kriegeskorte N (2015) Visual representations are dominated by intrinsic fluctuations correlated between areas. NeuroImage 114:275–286.

Henson RN, Goshen-Gottstein Y, Ganel T, Otten LJ, Quayle A, Rugg MD (2003) Electrophysiological and Haemodynamic Correlates of Face Perception, Recognition and Priming. Cereb Cortex 13:793–805.

Henson RN, Mouchlianitis E, Friston KJ (2009) MEG and EEG data fusion: Simultaneous localisation of face-evoked responses. NeuroImage 47:581–589.

Huang M-X, Song T, Hagler DJ Jr., Podgorny I, Jousmaki V, Cui L, Gaa K, Harrington DL, Dale AM, Lee RR, Elman J, Halgren E (2007) A novel integrated MEG and EEG analysis method for dipolar sources. NeuroImage 37:731–748.

Isik L, Meyers EM, Leibo JZ, Poggio TA (2014) The dynamics of invariant object recognition in the human visual system. J Neurophysiol 111:91–102.

Kaneshiro B, Perreau Guimaraes M, Kim H-S, Norcia AM, Suppes P (2015) A Representational Similarity Analysis of the Dynamics of Object Processing Using Single-Trial EEG Classification. PLoS ONE 10:095620.

Kiani R, Esteky H, Mirpour K, Tanaka K (2007) Object Category Structure in Response Patterns of Neuronal Population in Monkey Inferior Temporal Cortex. J Neurophysiol 97:4296–4309.

Kietzmann TC, Gert AL, Tong F, König P (2016) Representational Dynamics of Facial Viewpoint Encoding. J Cogn Neurosci:1–15.

King J-R, Dehaene S (2014) Characterizing the dynamics of mental representations: the temporal generalization method. Trends Cogn Sci 18:203–210.

Kriegeskorte N (2008) Representational similarity analysis – connecting the branches of systems neuroscience. Front Syst Neurosci 2:4.

Kriegeskorte N, Kievit RA (2013a) Representational geometry: integrating cognition, computation, and the brain. Trends Cogn Sci 17:401–412.

Kriegeskorte N, Kievit RA (2013b) Representational geometry: integrating cognition, computation, and the brain. Trends Cogn Sci 17:401–412.

Kriegeskorte N, Mur M, Ruff DA, Kiani R, Bodurka J, Esteky H, Tanaka K, Bandettini PA (2008) Matching Categorical Object Representations in Inferior Temporal Cortex of Man and Monkey. Neuron 60:1126–1141.

Kurth-Nelson Z, Barnes G, Sejdinovic D, Dolan R, Dayan P (2015) Temporal structure in associative retrieval. eLife 4:095620.

Kurth-Nelson Z, Economides M, Dolan RJ, Dayan P (2016) Fast Sequences of Non-spatial State Representations in Humans. Neuron 91:194–204.

Leahy RM, Mosher JC, Spencer ME, Huang MX, Lewine JD (1998) A study of dipole localization accuracy for MEG and EEG using a human skull phantom. Electroencephalogr Clin Neurophysiol 107:159–173.

Liu AK, Dale AM, Belliveau JW (2002) Monte Carlo simulation studies of EEG and MEG localization accuracy. Hum Brain Mapp 16:47–62.

Liu T, Slotnick SD, Serences JT, Yantis S (2003) Cortical Mechanisms of Feature-based Attentional Control. Cereb Cortex 13:1334–1343.

Maldjian JA, Laurienti PJ, Kraft RA, Burdette JH (2003) An automated method for neuroanatomic and cytoarchitectonic atlas-based interrogation of fMRI data sets. NeuroImage 19:1233–1239.

Maris E, Oostenveld R (2007) Nonparametric statistical testing of EEG- and MEG-data. J Neurosci Methods 164:177–190.

Marti S, King J-R, Dehaene S (2015) Time-Resolved Decoding of Two Processing Chains during Dual-Task Interference. Neuron 88:1297–1307.

Molins A, Stufflebeam SM, Brown EN, Hämäläinen MS (2008) Quantification of the benefit from integrating MEG and EEG data in minimum ℓ2-norm estimation. NeuroImage 42:1069–1077.

Müller K, Mika S, Rätsch G, Tsuda K, Schölkopf B (2001) An introduction to kernel-based learning algorithms. IEEE Trans Neural Netw 12:181–201.

Nichols TE, Holmes AP (2002) Nonparametric permutation tests for functional neuroimaging: A primer with examples. Hum Brain Mapp 15:1–25.

Pantazis D, Nichols TE, Baillet S, Leahy RM (2005) A comparison of random field theory and permutation methods for the statistical analysis of MEG data. NeuroImage 25:383–394.

Pflieger ME, Simpson GV, Ahlfors SP, Ilmoniemi RJ (2000) Superadditive Information from Simultaneous MEG/EEG Data. In: Biomag 96 (Aine CJ, Stroink G, Wood CC, Okada Y, Swithenby SJ, eds), pp 1154–1157. Springer New York.

Sharon D, Hämäläinen MS, Tootell RBH, Halgren E, Belliveau JW (2007) The advantage of combining MEG and EEG: Comparison to fMRI in focally stimulated visual cortex. NeuroImage 36:1225–1235.

Tadel F, Baillet S, Mosher JC, Pantazis D, Leahy RM (2011) Brainstorm: A User-Friendly Application for MEG/EEG Analysis. Comput Intell Neurosci 2011:1–13.

Taulu S, Kajola M, Simola J (2004) Suppression of interference and artifacts by the Signal Space Separation Method. Brain Topogr 16:269–275.

Taulu S, Simola J (2006) Spatiotemporal signal space separation method for rejecting nearby interference in MEG measurements. Phys Med Biol 51:1759.

Yoshinaga H, Nakahori T, Ohtsuka Y, Oka E, Kitamura Y, Kiriyama H, Kinugasa K, Miyamoto K, Hoshida T (2002) Benefit of Simultaneous Recording of EEG and MEG in Dipole Localization. Epilepsia 43:924–928.

